# Effects of Bimodal Olfactory and Mechanosensory Inputs in the Antennal Lobe of the Honeybee *Apis mellifera*

**DOI:** 10.64898/2026.04.08.717065

**Authors:** Shawn Mahoney, Shruti Joshi, Brian Smith, Mainak Patel, Hong Lei

## Abstract

Animals often aggregate information from multiple different sensory modalities to accurately assess and react to a stimulus. It is often assumed that cross-modality integration mostly occurs at high-level processing centers, such as the mammalian cortex or insect mushroom bodies. However, we hypothesized that integration could occur relatively early in the sensory pathways. The insect antennal lobe is one such location, receiving direct inputs from the antennae via the antennal nerve. These inputs are highly multimodal, including olfactory, mechanosesnory, and gustatory information, all of which are relevant to foraging honeybees (*Apis mellifera*). Here we assess integration by recording electrophysiological spike data within the honeybee antennal lobe while exposing the bee to various combinations of wind speed and odor concentration. This paper accompanies another publication by Joseph Reed and Mainak Patel approaching the same question from a modelling perspective, where their model corroborates our data and vice versa. Together, we show that integration occurs within this early layer of processing, while also demonstrating the complex relationship of these two closely-linked stimuli.

**SIGNIFICANCE:** Accurate perception depends on the brain’s ability to combine information from multiple senses, commonly thought belonging to high level of information processing. Using the European honeybee, *Apis mellifera*, we show strong evidence that olfactory and mechanosensory signals interact at an early stage of neural processing, within the antennal lobe, producing stimulus representations closely-linked to the animal’s navigation and decision-making. By identifying a tractable model for early multisensory processing, this work offers broader insight into how animals construct reliable representations of their environment.

## INTRODUCTION

Multimodal sensory integration creates holistic representations about stimuli so that animals can react appropriately. This can occur on many levels, from a single sensillum to the central layers of neural processing (Getz & Akers, 1994; Lei et al., 2022). Thus, characterizing this integration can be difficult, as the role of both each individual modality must be understood along with their interactions (Koziol et al., 2011; Mesulam, 1998; Verhagen & Engelen, 2006). For example, olfaction and mechanosensation are intrinsically linked because there are two components to an olfactory stimulus: The odorant molecules which bind to olfactory receptors, and the mechanosensory input of the air currents carrying those molecules (Goulet et al., 2025; Jernigan et al., 2026).

One common assumption is that olfactory processing displays greater integration at higher levels of neural processing like the mammalian piriform cortex, and less at lower levels (Emery et al., 2022; Verhagen & Engelen, 2006). However, evidence suggests that integration can occur much earlier in the processing pathway (Getz & Akers, 1994; Tiraboschi et al., 2021). In mice, the olfactory epithelium can respond to mechanical stimulation due to activation of G-protein-coupled receptors (Connelly et al., 2015; Grosmaitre et al., 2007). The insect antennal lobe (AL) presents an opportunity for studying integration at an early stage of processing, receiving direct inputs from peripheral sensory structures on the antenna via the antennal nerve, which carries axons from sensory neurons of various modalities distributed along the antenna (Lee & Strausfeld, 1990). AL neurons can be broadly categorized into two types. Projection Neurons (PNs), analogous to mammalian mitral/tufted cells, transmit information between the AL and other brain regions (Menzel et al., 2005; Mori & Shepherd, 1994), while Local Interneurons (LNs) process information within the AL (Fonta et al., 1993; Galizia & Kimmerle, 2004; Lei et al., 2002). LNs also modulate PN activity via the release of inhibitory neuromodulators like GABA (Girardin et al., 2013; Lei et al., 2002; Mustard et al., 2020; Sachse & Galizia, 2002).

The honeybee (*Apis mellifera*) is ideal for studying multimodal olfactory integration because of its well-characterized olfactory system, their antennae being highly multimodal organs capable of detecting both olfactory and mechanosensory inputs (de Brito Sanchez, 2011; Dreller & Kirchner, 1993; Haase et al., 2011; Scheiner et al., 2005). This relationship is ethologically relevant, as odor molecules are carried to the antennae by air flow from wind or the bee’s relative motion, which they use to navigate odor plumes toward flowers when foraging. This implies that olfactory inputs are relative to the speed and direction of wind over the antennae, and these inputs are sensitive to environmental wind fluctuations (Schneider et al., 1998). In such a dynamic sensory environment, bees must react quickly to changes in both stimuli.

There are two competing hypotheses for how integration might occur. First, information is integrated through separate channels, meaning the combined response is greater in intensity with the two modalities augmenting each other. Second, the two modalities could come through the same channel, meaning the interaction occurs prior to arrival at the AL. While there may be similar effects of each stimulus individually, the combined response would be different from the individual effects of each modality. This could be as simple as a multiplicative effect, or it could result in a different response entirely.

Here, we assess each of these hypotheses by exposing honeybees to various combinations of air speed and odor concentration while monitoring neural responses using extracellular spike recordings within the AL. We categorize the integration within PNs and LNs using the latency, firing rate, and general shape of the response. Our empirical data complement an accompanying paper, where (Reed & Patel, accompanying submission) independently created a mathematical model showing the integrative process within a network of PNs and LNs. Thus, we show that bimodal integration occurs on an earlier level of processing than initially assumed, and that there is a complex temporal interaction throughout the process.

## MATERIALS & METHODS

### Animals Used

We used the domestic European honeybee, *Apis mellifera*, sourced from the cordovan strain from Pendelton Apiaries in Stonyford, California. Bees were maintained outdoors in small colonies from which outgoing foragers were collected in vials and brought inside for daily experiments.

### Dissection and Recording Procedures

Bees were dissected in accordance with standard procedures (Lei et al., 2022). The bees were placed upon a 3D-printed stage, whereupon their heads were secured from the back using low-melting-temperature dental wax, and their antennae immobilized in a W-shaped position with eicosane to prevent interference in the dissection and recording processes. Once immobile, the head cap was removed and the AL uncovered. A NeuroNexus® acute probe with 16 contact sites was placed in the AL, and Kwiksil® tissue-compatible non-toxic adhesive was applied to the opening to form a cap, preventing desiccation and stabilizing the recording.

For each recording, bees were exposed to various odor-air speed combinations while we simultaneously recorded extracellular spiking activity within the AL. Recordings took 3-4 hours, with stimulus presentations divided into schemes based on air speed and odor concentration. Air was puffed for 0.5 seconds, with a 10-second waiting period between presentations to prevent adaptation. Thresholds for spike detection were set manually using visual analysis, with the general goal of capturing all spikes above 95% of noise. This approach allowed us to isolate AL neural activity, and any remaining noise artifacts could be manually sorted out afterwards using the Plexon® Offline Sorter© program.

### Air Speeds and Odor Concentrations

We generated variation in air speed using an adjustable olfactometer connected to an air supply. Speeds were created along an exponential gradient by combining multiple different airflow streams, resulting in 8 speeds: 0.34, 0.63, 1.1, 2.1, 4.2, 6.3, 12.6, and 25.3 m/s. Bees fly at an average of 7.5 m/s, topping out at 9 m/s maximum speed, meaning that first seven speeds are likely within a range the bee would naturally experience (Wenner, 1963). The last (25.3 m/s) wind speed is outside the natural range. Similarly, we used six odor concentrations, represented as the ratio of odor to mineral oil solvent: Control (no odor), 1:100, 1:10, 1:1, 10:1, and 100:1. Thus, there were a total of 48 air speed-odor combinations.

### Spike Sorting Analysis

After each recording, spiking data was sorted using the Offline Sorter program, and the sorted spikes were visualized for responsiveness using peristimulus raster and histogram plots generated within the NeuroExplorer© program. Any unit with a peristimulus spiking rate that noticeably rose above the baseline firing rate was considered to have responded to the stimulus. Units were only required to respond to the highest stimulus intensities to be categorized as “responsive”. From there, spikes were further sorted into projection neurons (PNs) and local interneurons (LNs) using k-means clustering based on spiking properties such as the spontaneous firing rate, interspike interval, and response “burstiness”, as outlined in (Lei et al., 2022; Meyer et al., 2013).

### PN/LN Separation

We identified a total of 34 PNs and 156 LNs, as well as 20 units which could not be sorted. This ratio of roughly 1 PN to 4.5 LNs corresponds to the 1:4 ratio described in (Meyer et al., 2013). We tested the strength of the separation by comparing the putative PNs and LNs using multiple methods. Across the board, the PNs had a higher firing rate than LNs, both when comparing their spontaneous firing rates (23.76 vs 4.24 spikes/s; t = 8.77, 188 df, p < 0.01), as well as their firing rate during stimulation (72.72 vs 15.86 spikes/s; t = 11.16, 188 df, p < 0.01). This means that there was an average increase from baseline of 48.96 spikes/s for PNs, compared with only 11.62 spikes/s for LNs (t = 11.29, 188 df, p < 0.01). PNs also had lower interspike intervals (0.024 vs 0.094 spikes/s; t = -7.71, 188 df, p < 0.01), with a higher average coefficient of variation (1.18 vs 1.06; t = 3.05, 188 df, p < 0.01), meaning that the PN spiking pattern was more “bursty”, or prone to clumping, while the LN pattern was closer to a random Poisson distribution. These statistics are all consistent with previous research, indicating that the PNs and LNs were accurately separated (Meyer et al., 2013). Supplemental Figure 1 shows examples of the separations.

### Response Measures and Data Analysis

We performed several analyses on sorted spikes to test for the integration of olfactory and mechanosensory inputs. The response window included the half-second stimulus presentation, as well as 1.5 seconds after the stimulus’s end, to account for both the bulk of the response, which can linger after the stimulus’s cessation, as well as “off” responses to the cessation itself. For the pre-stimulus comparison, we used spontaneous firing data from the 2-second window before the stimulus was given. From there, the response window was subdivided into 80 bins of 0.05 seconds each, with stimulus presentation occurring between bins 41-50.

Our primary analyses were based on two response parameters: The latency to start, and the response magnitude, defined using the net firing rate. The response start was defined as when the spiking rate rose more than 1 standard deviation above the resting mean following the stimulus onset at 0s. Two important notes are that the air flowed through a singular tube at constant pressure, and the bee antenna was positioned as close to the tube’s opening as possible without physically touching the tube (< 1 cm). Thus, the response latencies were minimally affected by the gap between the glass tube opening and the antenna. The net firing rate was calculated by taking the peristimulus rate throughout the 2-second response window and subtracting the spontaneous firing rate before the stimulus presentation.

Each of these parameters were assessed using a generalized linear mixed-effects model, with wind speed, odor concentration, and their interaction as fixed effects, and the individual neuronal units nested within each bee as a random effect. We generated both separate models for PNs and LNs, as well as a combined model with PN vs LN, plus the interactions with air speed and odor concentration, as fixed effects. The separate models approach produced the best fit, so that is the approach we will report here, with PNs and LNs as their own model each. For the response latency, we tried both a Poisson distribution and a gamma distribution, both using a log link, on the bin number corresponding to the response start. We found that Poisson fit the PNs best, while gamma fit the LNs, though neither distribution changed our significance values qualitatively. For the response magnitude, we square-root transformed both the spontaneous and the peristimulus (not net) firing rate data to achieve a normal distribution. Our response variable was then the transformed peristimulus rate, while we added the spontaneous rate as a fourth fixed effect to account for the heavy correlation between the two measures. See Supplemental Table 1 for a full list of model analyses and results.

In addition to the previous analyses, we also categorized responses based on their shape independent of the overall firing rate. To that end, we used a truncated suite of air speeds (0.34, 2.1, 6.3, and 25.3 m/s), increasing the number of repetitions per air speed from 1 to 10 for each bee, for greater confidence in the shapes. We then designed a hierarchical clustering algorithm that used the normalized spiking data to create a dissimilarity matrix based on the Hausdorff distance between each individual presentation curve, and we sorted each curve using a maximum distance clustering algorithm, creating 4 generalized response shapes for both PNs and LNs. We checked the sorting by generating dendrograms with labeled clusters, which can be viewed in Supplemental Figure 2. Since these clusters are categorical data, we calculated their statistical significance using ordinal logistic regression.

Finally, we analyzed the probability of changing response types based on stimulus inputs. To do this, we used the variance in response types for each neuron to calculate the probability that the response shape will change given an increase in stimulus intensity. This was broken down by both air speed and odor concentration, such that we found the probability to change response shape at a constant air speed given an increase in odor concentration, and vice versa. For example, we might look at the probability of changing from Type 1 to Type 2 at a constant speed of 0.34 m/s, given an increase in odor concentration. This gives us a metric to show the complex temporal dynamics involved when integrating bimodal sensory information.

## RESULTS

Neurons can encode information either by temporal coding, in which the latency or precise timing of spikes carries stimulus information, or by rate coding, in which information is represented mainly by the number of spikes fired over a time window (Panzeri et al., 2010). To assess the effect of bimodal integration on neural responses in the antennal lobe, we quantified the responses of PNs and LNs using their response latency and firing rate in all pairwise combinations of 5 concentrations and 8 wind speeds.

### Response Latency

The latencies in PNs to the onset of responses to unscented to air decreased with increasing wind speed by an average of 88% from the lowest to highest speed (Fig.1A), indicating there is a strong mechanical effect on PN activity (GLMM, β = -0.15, SE = 0.013, t = - 12.17, p < 0.01). The trend of response latencies to increasing wind speed in LNs was less steep (GLMM, β = -0.05, SE = 0.0053, t = -9.47, p < 0.01), though it still decreased by 38% from lowest to highest speed (Fig.1B). There is also a strong fixed effect of odor concentration (GLMM, PNs: β = -0.32, SE = 0.018, t = -17.89, p < 0.01; LNs: β = -0.19, SE = 0.0075, t = - 25.51, p < 0.01). Furthermore, the relationship between wind speed and latency was altered by adding odor stimulation, suggesting an interaction between the two stimuli (GLMM, PNs: β = 0.027, SE = 0.0037, t = 7.2, p < 0.01; LNs: β = 0.0087, SE = 0.0015, t = 5.82, p < 0.05) (Fig.1C, D). When comparing the two natural speeds (6.3 & 12.6 m/s), the higher wind speed produced shorter latency (darker grey bars, Fig.1C) as expected, but this trend was only maintained at the control and the lowest concentration (1:100). At higher concentrations, this trend was no longer apparent, suggesting a more complex bimodal integration. LNs did not show a consistent pattern across concentrations (Fig.1D).

**Figure 1:**
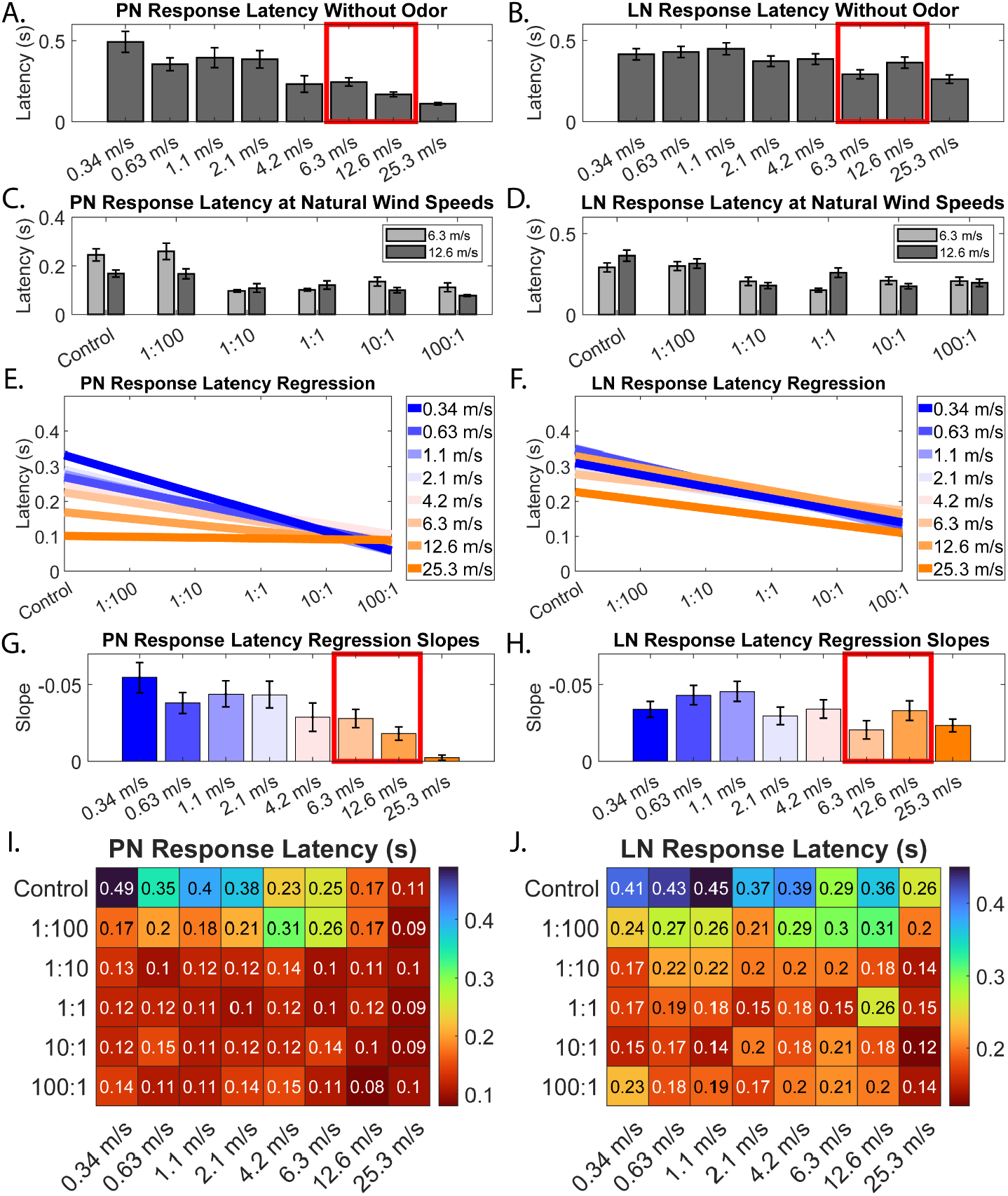
Response Latency, calculated as the time it takes for the spiking rate to rise 1 standard deviation above the resting rate. A, B) Change in latency at each unscented air speed in PNs (A) and LNs (B). Ecologically-relevant flying speeds are marked in the red box (Wenner, 1963). C, D) Bar graphs showing latency across odor concentrations at 6.3 and 12.6 m/s, which bracket the bee’s natural flying speed. E, F) Multiple linear regression plots showing the interaction between air speed and odor concentration in PNs (E) and LNs (F). Each line represents an air speed, showing the change in latency at that speed as concentration increases on the x-axis. Separation between the lines shows the effect of air speed, their slopes show the effect of odor, and the change in slope across lines shows the interaction. G, H) The slopes of the lines displayed in E and F, showing the slopes changing for each line. I, J) Heatmaps showing the latency in PNs (I) and LNs (J) under various wind speeds and odor concentrations.

To further reveal the effect of wind speed on odor responses, the response latencies were plotted against the whole range of concentrations as linear regressions (Fig.1E). Each line was derived from a particular wind speed across the range of odor concentrations. First, response latencies generally decreased with increasing wind speed, i.e. orange line (25.3 m/s) is at the lowest position and blue line (0.34 m/s) is at the highest position and other wind speeds are sequentially situated between these two lines. This order of response latencies was generally maintained at the concentration range, but the differences among the latencies (i.e. gap between the lines) shrank with increasing concentration, clearly showing the effect of odor stimulation on wind-induced responses. The stronger the odor stimulation, the samller the differences in response latency. Second, the slope of the regression lines became flatter systematically with increasing wind speed, indicating a weaker concentration dependency at higher wind speed. In other words, concentrations became less discernable when odors were delivered at high wind speed (Fig.1G). Together, there emerge a pattern where discriminability of the strength of mechanosensory and olfactory stimulation is downgraded by bimodal integration. The effect of bimodal integration was less clear in LNs (Fig.1F, H), as evident in all regression lines maintaining their slopes across concentrations. This pattern indicates that LNs can discriminate concentration steps regardless of the wind speed. Fig.1I & J emphasize this pattern by showing the inverse relationship between air speed and latency without odor as before, an effect which largely abates as odor concentration increases.

### Response Magnitude

Interestingly, the measurement of net spiking rate showed a different pattern (Fig.2), compared with the latency measurement. First, the series of increasing wind speeds did not systematically evoke increasing spiking rate at each concentration in both PNs and LNs (Fig.2A, B), except for the highest wind speed, which produced the strongest response in nearly all concentrations. That is, the orange lines are not always above the blue lines or vice versa. Second, the spiking rate generally increased with increasing concentration in all wind speeds, including the highest wind speed, although its regression line is flatter than other lines. These two observations suggest that concentration is a dominant factor here. We can verify this with our linear model, where while both concentration and wind speed are statistically significant, odor concentration has a higher effect size (GLMM, PNs: β = 0.49, SE = 0.037, t = 13.02, p < 0.01; LNs: β = 0.26, SE = 0.012, t = 21.9, p < 0.01) versus wind speed (GLMM, PNs: β = 0.16, SE = 0.029, t = 5.56, p < 0.01; LNs: β = 0.057, SE = 0.009, t = 6.33, p < 0.01). Even if we remove the possible outlier of the 25.6 m/s speed, these results do not qualitatively change. This means that, despite having a statistically significant interaction (GLMM, PNs: β = -0.02, SE = 0.0074, t = -2.68, p < 0.05; LNs: β = -0.012, SE = 0.0023, t = -4.57, p < 0.01), overall, Fig.2A-D shows that the trend in the spiking rate is not as easily discernible as with latency, lacking the clear, systematic change in slope between the two stimuli.

**Figure 2:**
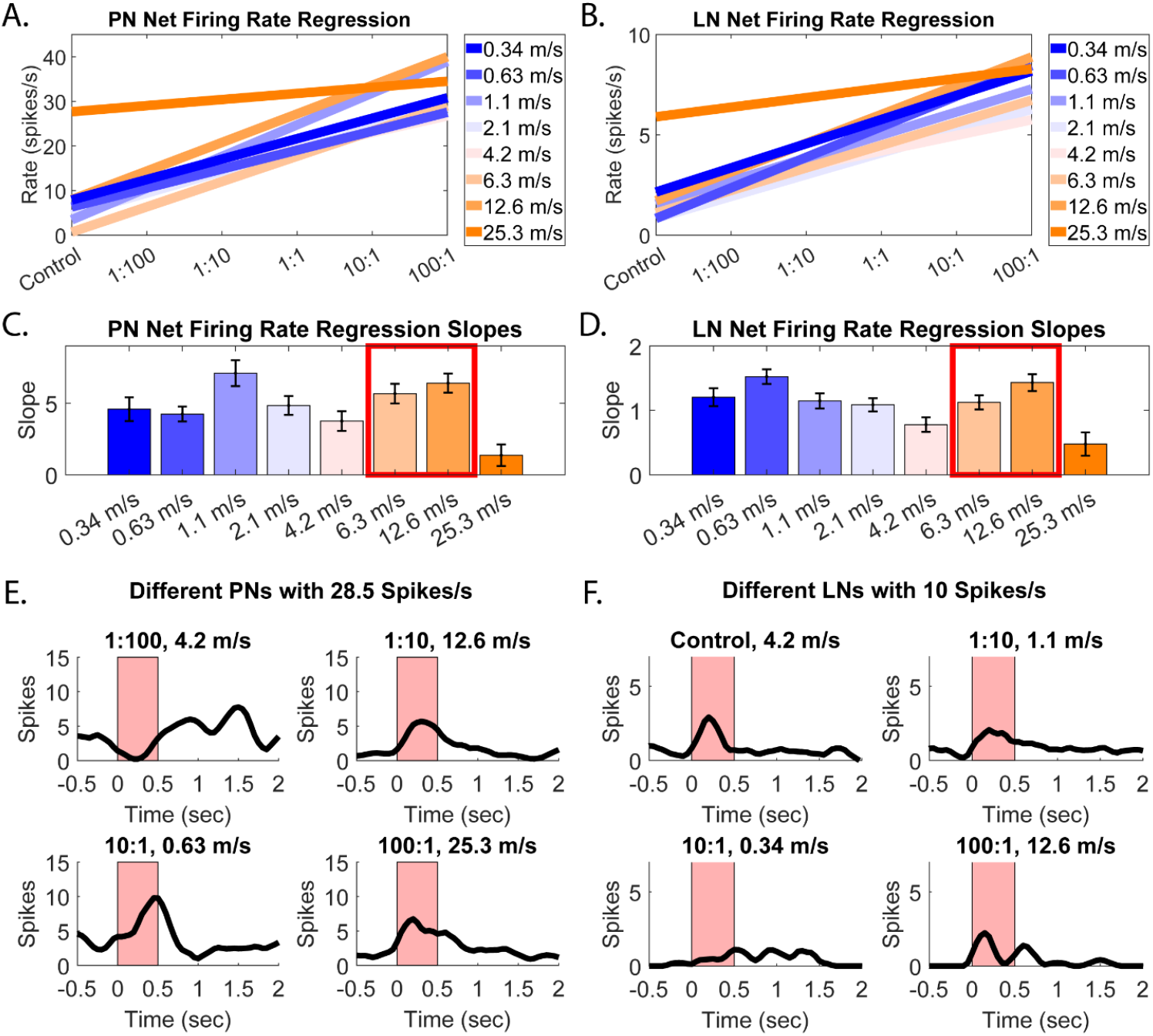
Firing rate over the entire 2-second response window. A, B) Multiple linear regression plots showing the interaction between air speed and odor concentration in PNs (A) and LNs (B). Each line represents an air speed, showing the change in firing rate at that speed as concentration increases on the x-axis. Separation between the lines denotes an effect of wind speed, their slope an effect of odor concentration, and change in slope between lines the interaction term. C, D) Bar plots showing the slopes of each line in A and B. Note the different y-axes between C and D. The gradual gradation seen in latency is not present here. E, F) Line plots showing 4 different PNs (E) which all have the same spiking rate, at different stimulus combinations. Similar plots are shown for LNs in (F). The firing rate over the entire response window is not an ideal measure because it does not account for temporal factors such as the size of the peak, the number of peaks, and so on.

The discrepancy between Fig.1 and Fig.2 prompted us to examine the peristimulus time histograms (PSTH) of responses. Fig.2E-F show that under different combinations of wind speed and concentration the specific shape can vary significantly. Thus, a better characterization of the interactions between wind speed and concentration would consider both the latency and temporal dynamics in spiking rate, i.e. the entire response shape.

### Response Shapes

We used a Hausdorff distance-based clustering algorithm to sort through all response curves (see Methods) and identified four response types for both PNs and LNs. Shapes for both types of neurons were similar, and there was no clear correlation between LN and PN response shapes, so for simplicity we only display results for the PNs here. Figure 3 displays the sorted response types for the PNs. Those types were:

**Figure 3:**
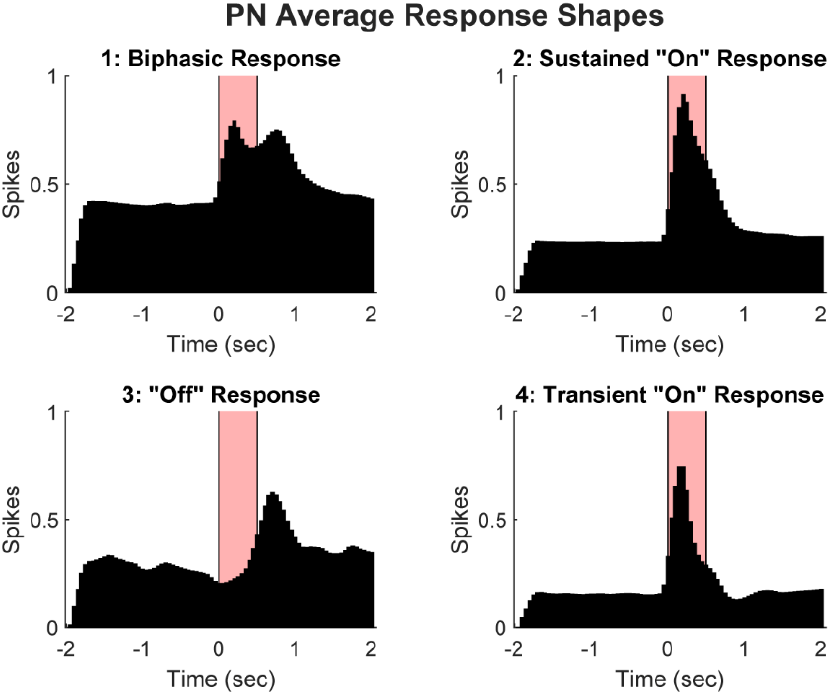
PN response shapes, as identified using a maximum clustering algorithm based off the Hausdorff distance between each individual response curve. In all, four response types were generated: 1. The biphasic response, characterized by two peaks, one at the beginning of stimulation and one at the end. 2. The sustained “on” response, which has a sharp peak at the beginning of the stimulus presentation and continues after presentation ends. 3. The “off” response, where the neuron only responds after the cessation of stimulus, often paired with inhibition during the stimulus presentation. 4. The transient “on” response, which like the sustained “on” response, peaks at the beginning of the stimulus, but ends quickly before the stimulus has finished.

1. Biphasic response: There are two peaks, one at the beginning of the stimulus presentation and one after termination.
2. Sustained “on” response: There is an increase in activity at the beginning of the stimulus presentation, which continues throughout stimulation and for some time after the presentation has ended.
3. “Off” response: Activity increases after the cessation of the stimulus. “Off” responses are usually weak and often reflect suppressed or inhibited firing during the stimulus presentation.
4. Transient “on” response: There is a transient peak with a short latency after the stimulus start, but activity ceases earlier, before or shortly after the end of the stimulus. Though this seems similar to Type 2, it is critical to note that these responses also show an over-suppression at the end of the response. Combining the transient and sustained “on” responses would also subsume the “off” response. See Supplemental Figure 2 for the dendrogram showing this separation.

To evaluate whether the response shapes were affected by wind speed and odor concentration, we placed the summed occurrences of reach response shape in a matrix with wind speeds in the horizontal dimension and concentrations at the vertical dimension (Fig.4). The relative height of bars represents the distribution of response shapes at a particular condition. Several features emerged from this analysis. First, among all four response shapes, Type 3 (“off” response) was the least frequent in all combinations. Second, wind speed seemed to be the dominating factor in shaping the response’s shape. In the non-scented control group, there was a relatively even distribution of response types at lower wind speeds, meaning that the response shape tended to be highly variable. As stimulus intensity increased to 25.3 m/s, the responses became heavily biased toward a sustained “on” response (Type 2), with a significant minority of transient “on” response (Type 4) as well, reflecting a decrease in latency and increase in overall length. This transition at the highest speed occurred at the other odor concentrations, confirming the dominant change brought by high wind speed.

**Figure 4:**
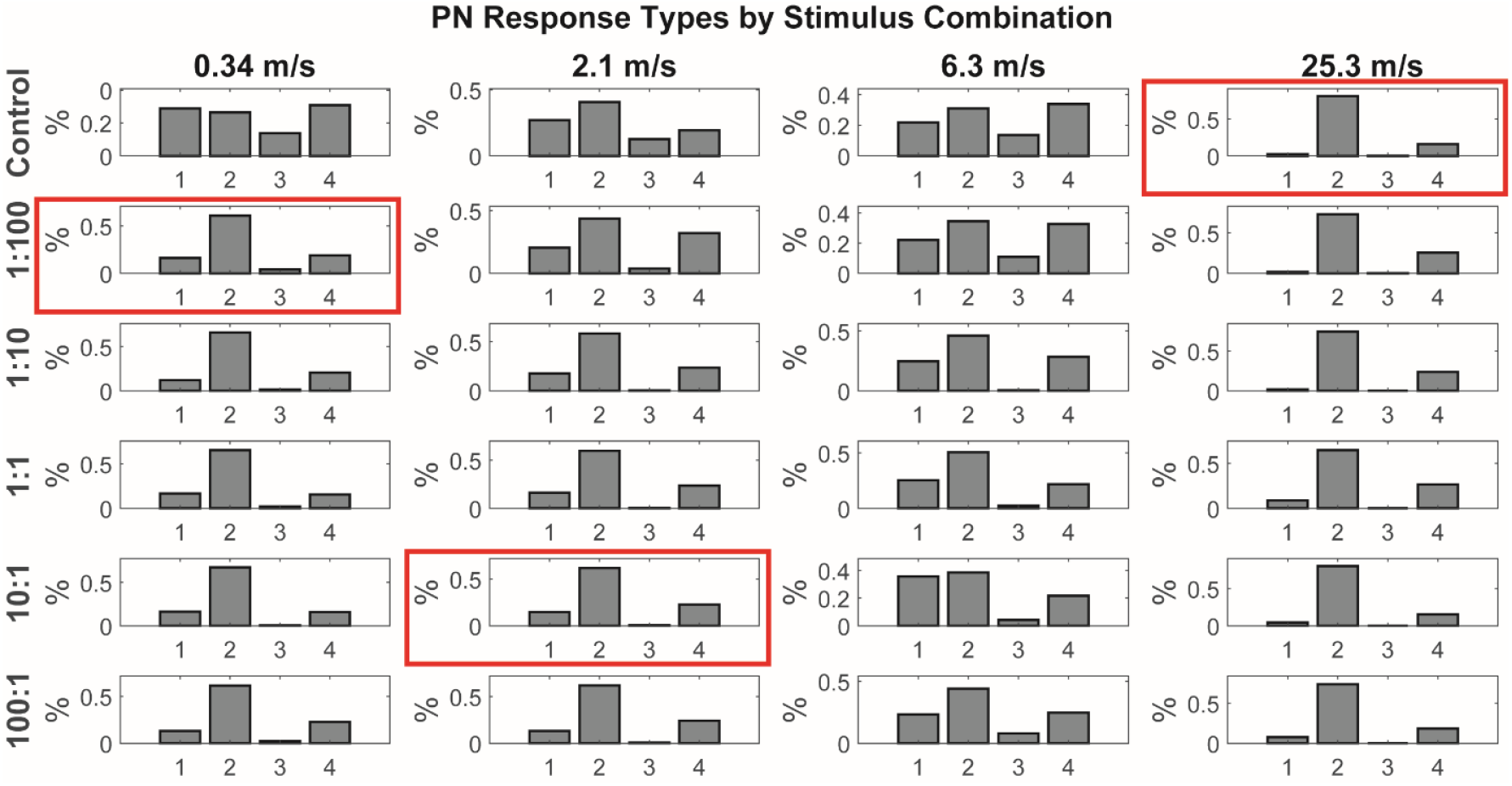
Proportions of response shapes in PNs by each speed-concentration pair. The y-axis scale for each pair is selected to best show the relative proportion of each response shape, and is thus different for each pair. At lower stimulus intensities, the responses are evenly distributed, but when traveling left to right or top to bottom, responses coalesce primarily around Type 2, the sustained “on” response, with a significant minority remaining in Type 4, the transient “on” response. Red boxes indicate the odor concentration at which Type 2 exceeds 60% of responses, for each wind speed. At 0.34 m/s, Type 2 exceeds 60% as soon as odor is added, at the 1:100 concentration. At 2.1 m/s, it takes much longer to dominate, reaching 60% at the 10:1 concentration. Type 2 never dominates at 6.3 m/s, and at 25.3 m/s, the responses saturate and Type 2 dominates throughout. Thus, odor concentration serves to homogenize responses, while wind speed serves to diversify them.

Third, and perhaps most interestingly, the dominant Type 2 response was not unique to high wind speed; odor stimuli produced a similar pattern even starting from the lowest concentration and slowest wind speed (1:100 onwards at 0.34 m/s). However, as shown by the red boxes in Figure 4, the odor-induced Type 2 dominance was weakened as wind speed increased until the highest wind speed, where, as seen in the other metrics, the responses saturate. These observations demonstrate that Type 2 responses could be activated by both wind and odor stimulations but at different intensities. Only the highest wind speed could achieve an equivalency of the lowest concentration. Below the highest wind speed, increasing wind speed tended to diversify response shapes by transforming the Type 2 response to other types. Correspondingly, increasing odor concentration tended to homogenize the response shapes around Type 2. The relative proportion of all four response shapes changed as the result of reducing the Type 2 response, raising an intriguing question about the mechanisms underlying the redistribution of response shapes.

### Response Type Changers

Since the proportion of response shape changes depended on the stimulus pairing, we determined the specific units that change response type, as well as the nature of the change. Figure 5 shows this using the probability of change when going from the 1:100 to 100:1 concentrations, at each wind speed. The control condition without odor was excluded, as we were more interested in the combined effect of wind speed and odor concentration, which unscented air does not show.

**Figure 5:**
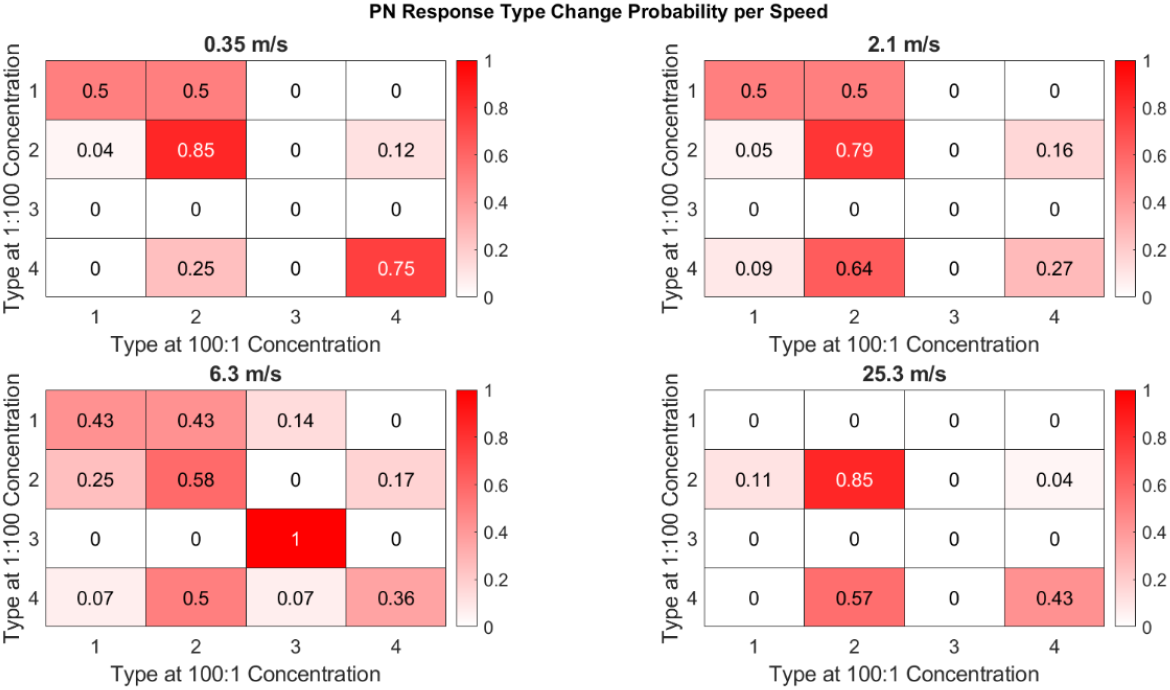
Probability to change response shape when going from 1:100 to 100:1 odor concentration at each wind speed. The y-axis represents the starting shapes at the 1:100 concentration, while the x-axis represents the ending shapes at the 100:1 concentration. Thus, the upper-left corner represents the probability that Type 1 responses remain at Type 1 when increasing the concentration, the box to the right represents the probability that they change to Type 2 when increasing the concentration, and so on. In other words, each heatmap is read row-by-row, moving left to right within each row.

Starting at the lowest air speed, 0.35 m/s, Figure 5 shows a strong coalescing effect upon Type 2. There is a 50% chance that Type 1 responses will change to Type 2, and an 85% chance that the already-dominant Type 2 responses will remain at Type 2 when increasing the odor concentration. By contrast, Type 4 responses have only a 25% chance of transitioning to Type 2, meaning that the majority of Type 4-responding neurons continue to do so regardless of odor concentration. Type 3 responses are present in such low proportion at this speed that they do not register one way or the other. This shows that Type 2 responses not only start in a dominant position, but remain dominant at this lowest wind speed.

Moving up to 2.1 m/s, we still see similar patterns, except now there is a higher propensity for Type 4 responses to transition to Type 2. Moving over to 6.3 m/s, however, we can see the true diversity of responses brought on by the air speed effect. We know that Type 2 never exceeds 60% of responses at 6.3 m/s (Fig.5), so while there is still a tendency to coalesce around Type 2, there is a higher likelihood for Type 1 or 4 responses to transition to a shape other than 2. Finally, when we look at the highest speed, 25.3 m/s, the responses saturate, and Type 2 dominates throughout, though with a significant minority which start in Type 4 and remain there across concentrations.

## DISCUSSION

### Integration Occurs Within the Antennal Lobe

Integration of olfactory and mechanosensory inputs occurs in early layers of the honeybee olfactory processing pathway. Consistent with literature (Lei et al., 2022; M. Patel et al., 2022; Tiraboschi et al., 2021; Tuckman et al., 2021), we report additional evidence for integration in PNs, showing a significant interaction between air speed and odor concentration in the latency to a sensory response. PN responses become stronger (i.e. shorter latency) with increasing air speed or odor concentration (Fig.1A, B). Higher odor concentrations shrank the latency differences among the wind speeds; likewise, higher wind speeds reduced sensitivity to differences among concentrations, as expressed by flatter regression slopes (Fig.1C, D). This phenomenon cannot be simply explained by a refractory mechanism because our experimental design allows us to observe gradual changes as these two factors affect each other (Fig.1C).

One implication of this integration is that a certain response strength can be achieved by various combinations of wind speed and concentration, as seen by imagining a horizontal line drawn through the regression lines in Fig.1C. In other words, a lower concentration at higher wind speed would achieve the same response as a higher concentration at lower wind speed, suggesting that these two modalities exert impacts on PNs inter-connectedly. Such inter-connectedness could be achieved by colocalization of receptors of different modalities on the same neuron, since the two receptors would then share the same pool of intracellular resources (i.e., calcium stores).

PNs receive direct input from sensory neurons on the antenna. One scenario is that bimodal integration occurs at the olfactory sensory neurons (OSNs), which then transmit those results to PNs. Though it is currently unclear whether the OSNs on honeybee antennae carry mechanical receptors, evidence from other animal models suggests this may be the case. In *Drosophila*, Or42b neurons, mostly responding to short-chain aliphatic esters, rely on the mechanosensitive ion channel TMEM63 to mediate humidity sensing (hygrosensation) (Li et al., 2022). Humidity changes cause the sensilla to bend, activating TMEM63. In zebra fish, mechanosensory proteins (Piezo 1 and Piezo 2) are found in various types of cells in the olfactory epithelium, including olfactory receptor cells (Aragona et al., 2024). In mice, the olfactory receptor proteins can be directly activated by mechanical stimuli (Connelly et al., 2015). Together, this points to the conclusion that OSNs are bimodal.

There is also evidence for integration in LNs, though notably weaker than in PNs (Fig.1B, D, F). When comparing the slopes between Fig.1G & H, we can see that, unlike the more even gradation observed in PNs, the decrease in slope for LNs, while still present, is flatter and less consistent. This could be, in part, because the LN network contains a broad range of neural morphologies (Fonta et al., 1993), meaning that averaging over all LNs results in more variation than in PNs.

### Integration Affects Response Shapes

While response latencies clearly show the effect of bimodal integration in PN responses, latency is not the only feature that postsynaptic neurons decode. The temporal dynamics within the entire response window can carry important stimulus information, demonstrated by many studies on spatiotemporal coding (Bathellier et al., 2010). When we examined the spiking rate within the entire 2-second response window (Fig.2), integrative patterns became more complex, which led us to analyze the response shapes in more detail.

We identified four response shapes, which show distinct temporal features (Fig.3). Both wind speed and odor concentration affect shapes in different ways. Non-scented air puffs produce a broad band of shapes across all speeds except for the non-natural highest speed (Fig.4). Adding odor homogenizes the shape around Type 2, the sustained “on” response, especially at the lowest wind speed. Thus, we can label Type 2 as the default response to odor stimulation. The Type 2 dominance gives way to other types of responses as wind speed increases (Fig.4, 5), until the highest speed. In other words, within a reasonable range, wind speed diversifies response shapes. The highest speed overloads the mechanosensory system, eliminating all response types but Type 2 regardless of concentration. Together, these results demonstrate (1) both types of stimuli affect response shapes; (2) the effects of bimodal integration over the full, 2-second time scale are non-linear and more complex compared to the shorter time scale (0.05s) measured by response latency.

Notably, the Type 3 “off” response always remains in low proportion, mostly disappearing with added odor concentration. The other three responses are all variations of “on” responses, which are consistent with the decreased latency from both air speed and odor concentration. Similarly, the odor-based coalescence on Type 2 responses is consistent with (Tuckman et al., 2021).

### Responses are Shaped by Excitatory and Inhibitory Inputs

The response shape is likely generated by a complex interaction between excitatory inputs and LN network inhibition (Cardin, 2018; M. J. Patel et al., 2013; Tuckman et al., 2021). As shown in (M. Patel et al., 2022), inhibition from the LNs slowly builds to diminish PN responses. During a sensory response, this slow inhibition from the LN network affects responses in both LNs and PNs. However, this inhibition is only produced during the actual stimulus window; after the stimulus turns off, so does the slow inhibition (Cardin, 2018), potentially resulting in a post-inhibitory rebound if there is lingering excitation.

Inhibition is likely a main driving factor behind the various response shapes seen in PNs. Take, for example, the Type 1 biphasic response. There is a strong excitatory push at the beginning, which slowly attenuates due to the buildup of inhibition over the stimulus window (Cardin, 2018; M. Patel et al., 2022; M. Patel & Joshi, 2015). Once the odor stimulus turns off, the inhibition ends (Cardin, 2018), resulting in a post-inhibitory bump in activity (Chang et al., 2018). This inhibition can also be overcome by sufficient excitatory activity (Moore et al., 2018), resulting in a Type 2 sustained “on” response, or if the excitatory response is short enough, the inhibition could be a non-factor, as in the Type 4 transient “on” response. The previously-described diversifying effect of wind speed serves to generate more Type 1 and 4 responses, both of which are more heavily affected by the buildup of slow inhibition. Therefore, it is plausible that extended mechanosensory inputs increase the buildup of slow inhibition within the AL, and that this buildup may account for the presence of the observed response shapes. In an accompanying publication, (Reed & Patel, accompanying submission) also demonstrate this effect using a model that simulates the interplay between excitation and slow inhibition, producing the same four response shapes that we see here.

### Response Shape is Adaptive in the Honeybee’s Odor Environment

Foraging honeybees operate in two contrasting olfactory environments. Inside the hive, waggle dances provide spatial information about food source direction and distance, which can then be combined with odor cues during the outbound search. At a distance from flowers, bees typically sample turbulent odor plumes composed of brief, intermittent filaments separated by clean air, making temporal structure a dominant feature of the signal (Conchou et al., 2019). By contrast, after entering a flower patch or returning to the hive, stimulation is prolonged and continuous, suggesting that honeybees may require distinct neural coding strategies for intermittent versus sustained odor input (Wechsler & Bhandawat, 2023). It’s plausible, then, that bees receive stronger mechanosensory stimulation during flight. Our data (Fig.4) show PNs produce more diversified response shapes under moderate wind speeds by weakening the dominant Type 2 response. Notably, the Type 4 transient “on” response emerges when wind speed increases. Such variable response shapes may provide advantages for tracking odor plumes during flight. In contrast, the Type 2 response may be optimal when bees are on a surface evaluating the surrounding odor environment. The channels sensing mechanical stimulation may function as an adjusting device on olfactory neurons to vary their response shapes.

### Integration is Strongest at the Bee’s Natural Flight Speed

We propose that the antennae may be tuned to integrate olfactory and mechanosensory stimuli at specific wind speeds. The strongest integration is seen in speeds that most closely mimic the bee’s natural flight speed. In the wild, a bee flies at an average of 7.5 m/s, up to a maximum of 9 m/s (Wenner, 1963), often following the odor plume upwind (Barron & Srinivasan, 2006), meaning the 6.3 and 12.6 m/s wind speeds bracket the bee’s natural range. These speeds are where we see the greatest integration, which saturates quickly with the addition of odor as shown in Figure 1C. At the 25.3 m/s speed, far outside the bee’s natural range (Wenner, 1963), all integration is completely overloaded by the mechanosensory effect of the wind. Figure 1E shows this, where the orange line representing the 25.3 m/s speed is flat, indicating little to no effect of odor at that highest speed. Thus, the bee is very sensitive to wind speeds between roughly 6-13 m/s, the speeds which it is most likely to experience when foraging, and too far above that results in overload.

## CONCLUSION

In this paper, we have demonstrated the bimodal integration of olfactory and mechanosensory stimuli within the honeybee AL. We show that there is an interaction in terms of the latency to the start of the response, as well as why whole-response measures like the firing rate may be not be ideal when analyzing responses to variation in multiple stimuli. We then show how the response shape is also affected by bimodal stimulation, where there is a disparate effect of olfaction and mechanosensation: Olfactory inputs serve to coalesce the responses toward faster and longer shapes, whereas mechanosensory inputs diversify the response shapes to include shorter or bimodal responses. This is further explored in the accompanying paper by (Reed & Patel, accompanying submission), where they explore the network dynamics underlying the generation of these different shapes. Together, these works serve to provide a model where integration occurs over diverse timescales, with the temporal pattern of the response being dependent on the specific properties of the stimuli being integrated.

## ACKNOWLEDGEMENTS

The authors would like to extend our sincerest thanks to everyone who has contributed to this paper, especially the scientists working within the ASU Social Insect Research Group and the cross-institution bimodality working group, whose continuous feedback has been instrumental in the writing of this paper. We’d like to particularly thank Ross Iaci, Stephen Pratt, and Theodore Pavlic for their contributions to our statistical analyses, as well as Abigail Brow and Kendrick Nakamura for consistently reviewing and providing feedback on this publication. We’d also like to acknowledge the accompanying publication, “Antennal Lobe Dynamics and the Generation of Diverse Response Patterns to Mechanosensory Stimulation”, as well as its authors Joseph Reed and Mainak Patel, for their continued collaboration on this project. Finally, we’d like to thank the NIDCD R01 DC020892-01 to Hong Lei and the NSF/CIHR/DFG/FRQ/UKRI-MRC Next Generation Networks for Neuroscience Program (Award #2014217) subaward to Brian Smith for their financial backing throughout this project.

---

**Figure S1:**
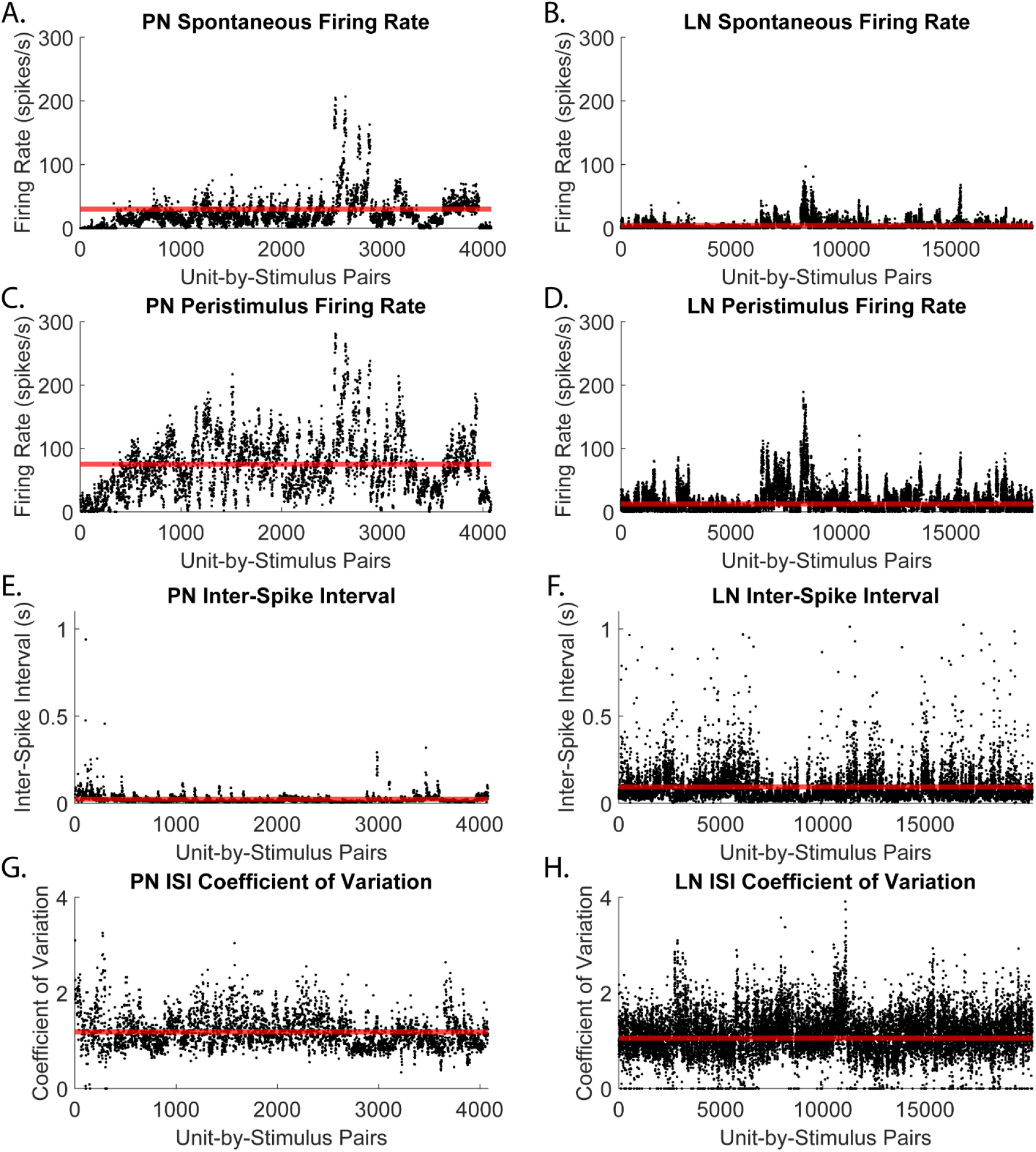
Comparisons between PNs and LNs. Red lines mark average values in each plot. A, B) Spontaneous firing rate before each stimulus presentation in PNs (A) and LNs (B). PNs have a higher average spontaneous firing rate than LNs. C, D) Peristimulus firing rate across all stimulus presentations in PNs (C) and LNs (D). The difference in firing rates is even greater when comparing the peristimulus firing rate with the spontaneous rate. E, F) Average peristimulus inter-spike intervals across all stimulus presentations in PNs (E) and LNs (F). LNs have higher inter-spike intervals on average than PNs. G, H) Coefficient of variation of inter-spike intervals across all stimulus presentations in PNs (G) and LNs (H). LN covariance is close to 1, indicating a Poisson distribution, whereas PN covariance is higher, meaning that the PN firing distribution is more “bursty”.

**Table S1:** Full results for the generalized linear mixed-effects models. For Latency, the formula was defined as: Latency ∼ 1 + concentration + speed + concentration:speed + (1|bee) + (1|bee:unit), using a Poisson distribution for the PNs and gamma for the LNs, both with a log link. For firing rate, the formula was defined as: Peristimulus rate ∼ 1 + spontaneous rate + concentration + speed + concentration:speed + (1|bee) + (1|bee:unit). Significant p-values are marked in bold.

|  |  | Latency |  |  |  |  |  |  |  |  |  |  |
| --- | --- | --- | --- | --- | --- | --- | --- | --- | --- | --- | --- | --- |
|  |  | β | Error | T-Stat | DF | p-Value | Lower CI | Upper CI | AIC | BIC | Log Likelihood | Deviance |
| PNs | Intercept | 2.427002 | 0.088686 | 27.3662 | 1592 | **1.84E-135** | 2.253048 | 2.600955 | 5163.661 | 5195.912 | -2575.830298 | 5151.661 |
|  | Concentration | -0.32079 | 0.017934 | -17.887 | 1592 | **2.40E-65** | -0.35597 | -0.28561 |  |  |  |  |
|  | Air Speed | -0.15455 | 0.012701 | -12.1688 | 1592 | **1.22E-32** | -0.17947 | -0.12964 |  |  |  |  |
|  | Concentration:Air Speed | 0.026948 | 0.003743 | 7.200107 | 1592 | **9.24E-13** | 0.019607 | 0.034289 |  |  |  |  |
| LNs | Intercept | 2.169694 | 0.101189 | 21.44193 | 6934 | **8.30E-99** | 1.971332 | 2.368056 | 20297.23 | 20345.14 | -10141.61498 | 20283.23 |
|  | Concentration | -0.18135 | 0.016179 | -11.2087 | 6934 | **6.55E-29** | -0.21307 | -0.14963 |  |  |  |  |
|  | Air Speed | -0.04828 | 0.012485 | -3.86724 | 6934 | **1.11E-04** | -0.07276 | -0.02381 |  |  |  |  |
|  | Concentration:Air Speed | 0.006423 | 0.003165 | 2.029258 | 6934 | **0.04247** | 0.000218 | 0.012627 |  |  |  |  |
|  |  | Firing Rate |  |  |  |  |  |  |  |  |  |  |
|  |  | β | Error | T-Stat | DF | p-Value | Lower CI | Upper CI | AIC | BIC | Log Likelihood | Deviance |
| PNs | Intercept | 0.80742 | 0.271423 | 2.974769 | 1627 | **0.002975** | 0.275045 | 1.339795 | 5229.921 | 5273.102 | -2606.960599 | 5213.921 |
|  | Concentration | 0.485407 | 0.037269 | 13.0245 | 1627 | **5.84E-37** | 0.412308 | 0.558507 |  |  |  |  |
|  | Air Speed | 0.159428 | 0.028684 | 5.558071 | 1627 | **3.18E-08** | 0.103167 | 0.21569 |  |  |  |  |
|  | Concentration:Air Speed | -0.01975 | 0.007382 | -2.67532 | 1627 | **7.54E-03** | -0.03423 | -0.00527 |  |  |  |  |
| LNs | Intercept | 0.919487 | 0.156878 | 5.861156 | 7483 | **4.79E-09** | 0.611962 | 1.227013 | 18196.63 | 18252 | -9090.314528 | 18180.63 |
|  | Concentration | 0.255201 | 0.011657 | 21.89253 | 7483 | **4.97E-103** | 0.23235 | 0.278052 |  |  |  |  |
|  | Air Speed | 0.056878 | 0.008988 | 6.328146 | 7483 | **2.62E-10** | 0.039259 | 0.074497 |  |  |  |  |
|  | Concentration:Air Speed | -0.01056 | 0.002308 | -4.57423 | 7483 | **4.86E-06** | -0.01508 | -0.00603 |  |  |  |  |

**Figure S2:**
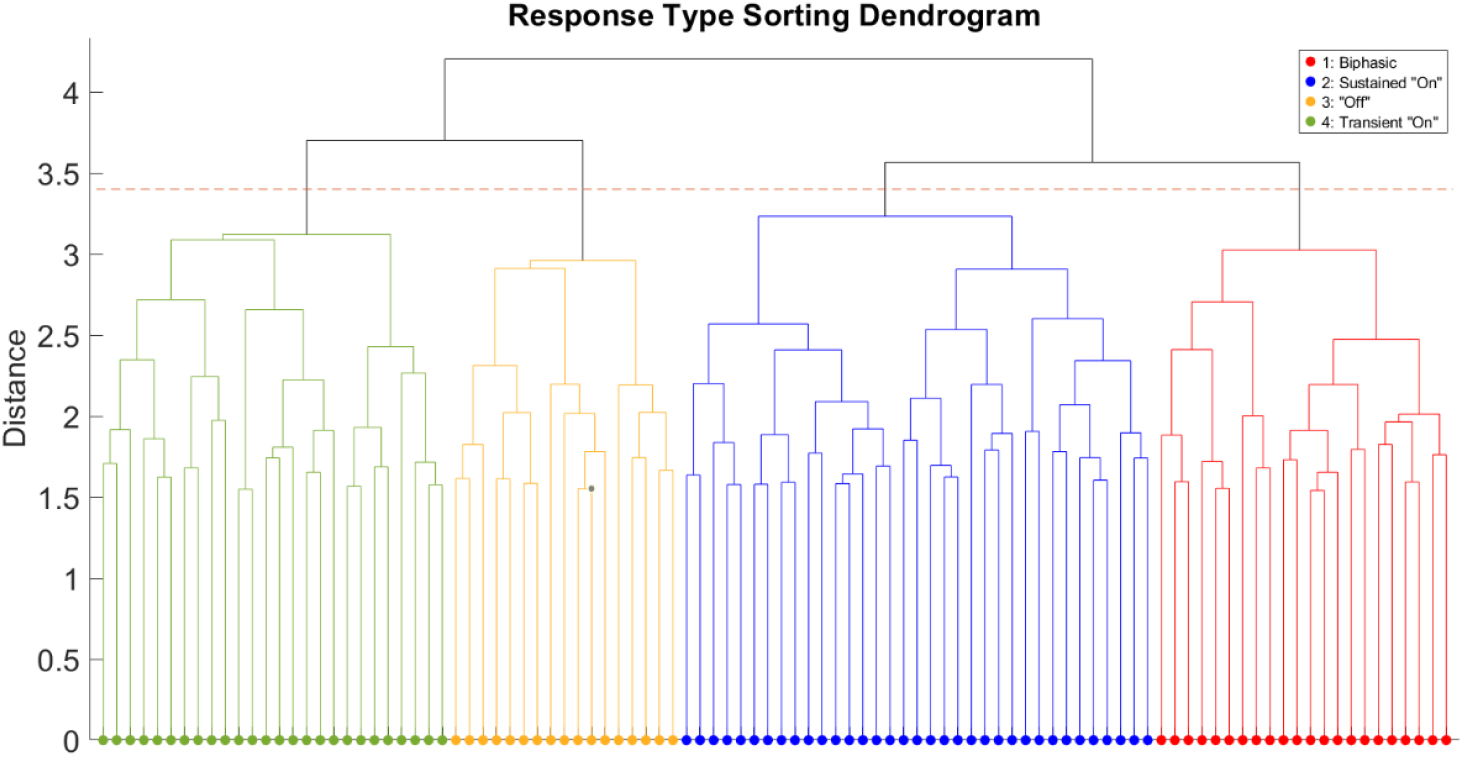
Dendrogram showing the sorting of response types. Colors represent the four distinct response shapes, and the red line indicates the distance cutoff. More similar shapes are grouped closer together, i.e. Types 1 (biphasic) is more similar to Type 2 (sustained “on”) than it is to Type 4 (transient “on”). The size of each color is proportional to the number of unit-stimulus pairs within each type category, meaning that Type 2 is the most numerous, followed by Type 4, Type 1, and Type 3.

## REFERENCES

Aragona, M., Mhalhel, K., Cometa, M., Franco, G. A., Montalbano, G., Guerrera, M. C., Levanti, M., Laurà, R., Abbate, F., Vega, J. A., & Germanà, A. (2024). Piezo 1 and Piezo 2 in the Chemosensory Organs of Zebrafish (Danio rerio). International Journal of Molecular Sciences, 25(13), 7404. 10.3390/ijms25137404

Barron, A., & Srinivasan, M. V. (2006). Visual regulation of ground speed and headwind compensation in freely flying honey bees (Apis mellifera L.). The Journal of Experimental Biology, 209(Pt 5), 978–984. 10.1242/jeb.02085

Bathellier, B., Gschwend, O., & Carleton, A. (2010). Temporal Coding in Olfaction. In A. Menini (Ed.), The Neurobiology of Olfaction. CRC Press/Taylor & Francis. http://www.ncbi.nlm.nih.gov/books/NBK55968/

Cardin, J. A. (2018). Inhibitory Interneurons Regulate Temporal Precision and Correlations in Cortical Circuits. Trends in Neurosciences, 41(10), 689–700. 10.1016/j.tins.2018.07.015

Chang, M., Dian, J. A., Dufour, S., Wang, L., Moradi Chameh, H., Ramani, M., Zhang, L., Carlen, P. L., Womelsdorf, T., & Valiante, T. A. (2018). Brief activation of GABAergic interneurons initiates the transition to ictal events through post-inhibitory rebound excitation. Neurobiology of Disease, 109, 102–116. 10.1016/j.nbd.2017.10.007

Conchou, L., Lucas, P., Meslin, C., Proffit, M., Staudt, M., & Renou, M. (2019). Insect Odorscapes: From Plant Volatiles to Natural Olfactory Scenes. Frontiers in Physiology, 10. 10.3389/fphys.2019.00972

Connelly, T., Yu, Y., Grosmaitre, X., Wang, J., Santarelli, L. C., Savigner, A., Qiao, X., Wang, Z., Storm, D. R., & Ma, M. (2015). G protein-coupled odorant receptors underlie mechanosensitivity in mammalian olfactory sensory neurons. Proceedings of the National Academy of Sciences, 112(2), 590–595. 10.1073/pnas.1418515112

de Brito Sanchez, M.G. (2011). Taste Perception in Honey Bees. Chemical Senses, 36(8), 675–692. 10.1093/chemse/bjr040

Dreller, C., & Kirchner, W. H. (1993). Hearing in honeybees: Localization of the auditory sense organ. Journal of Comparative Physiology A, 173(3), 275–279. 10.1007/BF00212691

Emery, B. A., Hu, X., Maugeri, L., Khanzada, S., Klutsch, D., Altuntac, E., & Amin, H. (2022). Large-scale Multimodal Recordings on a High-density Neurochip: Olfactory Bulb and Hippocampal Networks. Annual International Conference of the IEEE Engineering in Medicine and Biology Society. IEEE Engineering in Medicine and Biology Society. Annual International Conference, 2022, 3111–3114. 10.1109/EMBC48229.2022.9871961

Fonta, C., Sun, X.-J., & Masson, C. (1993). Morphology and spatial distribution of bee antennal lobe interneurones responsive to odours. Chemical Senses, 18(2), 101–119. 10.1093/chemse/18.2.101

Galizia, C. G., & Kimmerle, B. (2004). Physiological and morphological characterization of honeybee olfactory neurons combining electrophysiology, calcium imaging and confocal microscopy. Journal of Comparative Physiology. A, Neuroethology, Sensory, Neural, and Behavioral Physiology, 190(1), 21–38. 10.1007/s00359-003-0469-0

Getz, W. M., & Akers, R. P. (1994). Honeybee olfactory sensilla behave as integrated processing units. Behavioral and Neural Biology, 61(2), 191– 195. 10.1016/s0163-1047(05)80075-5

Girardin, C. C., Kreissl, S., & Galizia, C. G. (2013). Inhibitory connections in the honeybee antennal lobe are spatially patchy. Journal of Neurophysiology, 109(2), 332–343. 10.1152/jn.01085.2011

Goulet, D., Smith, B., True, A., & Crimaldi, J. (2025). Pore plate sensilla scale and distribution modulate odor capture around honey bee antennae. Scientific Reports, 15, 41574. 10.1038/s41598-025-25426-1

Grosmaitre, X., Santarelli, L. C., Tan, J., Luo, M., & Ma, M. (2007). Dual functions of mammalian olfactory sensory neurons as odor detectors and mechanical sensors. Nature Neuroscience, 10(3), 348–354. 10.1038/nn1856

Haase, A., Rigosi, E., Frasnelli, E., Trona, F., Tessarolo, F., Vinegoni, C., Anfora, G., Vallortigara, G., & Antolini, R. (2011). A multimodal approach for tracing lateralisation along the olfactory pathway in the honeybee through electrophysiological recordings, morpho-functional imaging, and behavioural studies. European Biophysics Journal, 40(11), 1247–1258. 10.1007/s00249-011-0748-6

Jernigan, C. M., Connor, E., Lei, H., Victor, J. D., Crimaldi, J., & Smith, B. H. (2026). Active sensing: Different plume structures affect movements of antennae in honey bees (Apis mellifera). Journal of Experimental Biology, 229(4), jeb250786. 10.1242/jeb.250786

Koziol, L. F., Budding, D. E., & Chidekel, D. (2011). Sensory Integration, Sensory Processing, and Sensory Modulation Disorders: Putative Functional Neuroanatomic Underpinnings. The Cerebellum, 10(4), 770–792. 10.1007/s12311-011-0288-8

Lee, J. K., & Strausfeld, N. J. (1990). Structure, distribution and number of surface sensilla and their receptor cells on the olfactory appendage of the male moth Manduca sexta. Journal of Neurocytology, 19(4), 519–538. 10.1007/BF01257241

Lei, H., Christensen, T. A., & Hildebrand, J. G. (2002). Local inhibition modulates odor-evoked synchronization of glomerulus-specific output neurons. Nature Neuroscience, 5(6), 557–565. 10.1038/nn0602-859

Lei, H., Haney, S., Jernigan, C. M., Guo, X., Cook, C. N., Bazhenov, M., & Smith, B. H. (2022). Novelty detection in early olfactory processing of the honey bee, Apis mellifera. PLOS ONE, 17(3), e0265009. 10.1371/journal.pone.0265009

Li, S., Li, B., Gao, L., Wang, J., & Yan, Z. (2022). Humidity response in Drosophila olfactory sensory neurons requires the mechanosensitive channel TMEM63. Nature Communications, 13(1), 3814. 10.1038/s41467-022-31253-z

Menzel, R., Galizia, G., Müller, D., & Szyszka, P. (2005). Odor Coding in Projection Neurons of the Honeybee Brain. Chemical Senses, 30(Suppl_1), i301–i302. 10.1093/chemse/bjh234

Mesulam, M. M. (1998). From sensation to cognition. Brain, 121(6), 1013–1052. 10.1093/brain/121.6.1013

Meyer, A., Galizia, C. G., & Nawrot, M. P. (2013). Local interneurons and projection neurons in the antennal lobe from a spiking point of view. Journal of Neurophysiology, 110(10), 2465–2474. 10.1152/jn.00260.2013

Moore, A. K., Weible, A. P., Balmer, T. S., Trussell, L. O., & Wehr, M. (2018). Rapid Rebalancing of Excitation and Inhibition by Cortical Circuitry. Neuron, 97(6), 1341-1355.e6. 10.1016/j.neuron.2018.01.045

Mori, K., & Shepherd, G. M. (1994). Emerging principles of molecular signal processing by mitral/tufted cells in the olfactory bulb. Seminars in Cell Biology, 5(1), 65–74. 10.1006/scel.1994.1009

Mustard, J. A., Jones, L., & Wright, G. A. (2020). GABA signaling affects motor function in the honey bee. Journal of Insect Physiology, 120, 103989. 10.1016/j.jinsphys.2019.103989

Panzeri, S., Brunel, N., Logothetis, N. K., & Kayser, C. (2010). Sensory neural codes using multiplexed temporal scales. Trends in Neurosciences, 33(3), 111–120. 10.1016/j.tins.2009.12.001

Patel, M. J., Rangan, A. V., & Cai, D. (2013). Coding of odors by temporal binding within a model network of the locust antennal lobe. Frontiers in Computational Neuroscience, 7. 10.3389/fncom.2013.00050

Patel, M., & Joshi, B. (2015). Modeling the evolving oscillatory dynamics of the rat locus coeruleus through early infancy. Brain Research, 1618, 181– 193. 10.1016/j.brainres.2015.05.033

Patel, M., Kulkarni, N., Lei, H. H., Lai, K., Nematova, O., Wei, K., & Lei, H. (2022). Experimental and theoretical probe on mechano- and chemosensory integration in the insect antennal lobe. Frontiers in Physiology, 13. 10.3389/fphys.2022.1004124

Reed, J., & Patel, M. (accompanying submission). Antennal Lobe Dynamics And The Generation Of Diverse Response Patterns To Mechanosensory Stimulation.

Sachse, S., & Galizia, C. G. (2002). Role of inhibition for temporal and spatial odor representation in olfactory output neurons: A calcium imaging study. Journal of Neurophysiology, 87(2), 1106–1117. 10.1152/jn.00325.2001

Scheiner, R., Schnitt, S., & Erber, J. (2005). The functions of antennal mechanoreceptors and antennal joints in tactile discrimination of the honeybee (Apis mellifera L.). Journal of Comparative Physiology A, 191(9), 857–864. 10.1007/s00359-005-0009-1

Schneider, R. W. S., Price, B. A., & Moore, P. A. (1998). Antennal morphology as a physical filter of olfaction: Temporal tuning of the antennae of the honeybee, Apis mellifera. Journal of Insect Physiology, 44(7), 677–684. 10.1016/S0022-1910(98)00025-0

Tiraboschi, E., Leonardelli, L., Segata, G., & Haase, A. (2021). Parallel Processing of Olfactory and Mechanosensory Information in the Honey Bee Antennal Lobe. Frontiers in Physiology, 12. 10.3389/fphys.2021.790453

Tuckman, H., Kim, J., Rangan, A., Lei, H., & Patel, M. (2021). Dynamics of sensory integration of olfactory and mechanical stimuli within the response patterns of moth antennal lobe neurons. Journal of Theoretical Biology, 509, 110510. 10.1016/j.jtbi.2020.110510

Verhagen, J. V., & Engelen, L. (2006). The neurocognitive bases of human multimodal food perception: Sensory integration. Neuroscience & Biobehavioral Reviews, 30(5), 613–650. 10.1016/j.neubiorev.2005.11.003

Wechsler, S. P., & Bhandawat, V. (2023). Behavioral algorithms and neural mechanisms underlying odor-modulated locomotion in insects. Journal of Experimental Biology, 226(1), jeb200261. 10.1242/jeb.200261

Wenner, A. M. (1963). The Flight Speed of Honeybees: A Quantitative Approach. Journal of Apicultural Research, 2(1), 25–32. 10.1080/00218839.1963.11100053

